# A Microneurosurgical Survival Platform for Elucidating Mechanisms of Brain Tumor Recurrence and Metastasis

**DOI:** 10.64898/2026.08.31.748338

**Authors:** Diganta Das, Brooke Nakamura, Josh Neman

## Abstract

Brain tumor recurrence remains the leading cause of mortality in neuro-oncology, and there is a lack of preclinical models replicating the clinical cycle of surgical resection and relapse. To bridge this gap, we developed a novel microneurosurgical survival platform in mice using the NICO Myriad system. We orthotopically implanted pediatric medulloblastoma cells into the mouse cerebral cortex or cerebellum, followed by longitudinal microneurosurgical resection. Bioluminescence imaging and gross fluorescence verified successful resection, local and distal recurrence and metastasis. Comparative bulk RNA sequencing revealed extensive stage-specific transcriptomic divergence alongside conserved core gene sets (2,702 genes in the cerebral cortex and 3,240 genes in the cerebellum) across primary, locally recurrent, and distally recurrent stages. Pathway analysis shows activation of cellular growth, second messenger signaling, and cellular stress adaptation pathways. Targeted qPCR validation demonstrated that post-surgical relapse is driven by a distinct molecular program: recurrent tumors downregulate primary developmental drivers (*PTCH1*, *MYCBP2*), canonical suppressors (*FOS*, *PTEN*), and chromatin regulators (*HDAC2*), while selectively upregulating post-transcriptional machinery (*RBM8A*), endosomal trafficking regulators (*RAB5C*), acetyltransferases (*NAA15*), and the m^6^A RNA demethylase *ALKBH5*. These findings reveal that medulloblastoma shifts from a primary oncogenic state toward post-transcriptional and transcriptomic survival mechanisms following surgery. Identifying persistent candidates within this conserved core framework provides a roadmap for next-generation precision immunotherapies.

## Introduction

The clinical management of brain tumor is complicated by molecular and phenotypic divergence that occurs between the primary tumor and its subsequent recurrence. Rather than a static progression, relapse is often characterized by a dynamic evolutionary event where the tumor undergoes extensive clonal selection and microenvironmental adaptation [1]. This transition often results in a mutator phenotype characterized by shifted epigenetic landscapes and altered metabolic demands that differ from the naive state [2]. This complexity is further exacerbated by the tumor’s ability to colonize anatomically distant niches such as the cerebellar cortex, where it survives under different selective pressures than those found in the native cerebellum [3, 4]. Understanding these mechanisms by which the tumor integrates into diverse brain environments requires a critical observation which requires high-fidelity longitudinal transcriptomic profiling that can capture the molecular changes in a clinically relevant manner.

Tumor resection is a technique that emphasizes surgical precision and prioritizes the preservation of the neurovascular architecture [5]. Clinical data demonstrate that an extent of resection (EOR) exceeding 98% of the contrast-enhancing volume is a primary predictor of overall survival (OS) in patients with glioblastoma (GBM) [6. However, despite the advancement of intraoperative adjuncts such as 5-aminolevulinic acid (5-ALA]. However, despite advances in intraoperative adjuncts such as 5-aminolevulinic acid, the surgical management of brain tumors remains complicated by the disease’s biological evolution. The relevance of microneurosurgical intervention shifts significantly between the primary and recurrent settings. In primary resections, surgeons contend with relatively naive tissue, whereas recurrent tumors exist within a distorted landscape of surgical scarring and radiation-induced necrosis, making the identification of the tumor-brain interface increasingly difficult [7]. A major limitation in current neuro-oncology is the inability to accurately study the variations between these states. While primary tumors are well-documented, how a tumor survives the selective pressure of chemotherapy and radiotherapy to emerge as a resistant recurrence remains poorly understood. This knowledge gap can be addressed by using *in vivo* models that replicate the clinical cycle of resection and recurrence.

Medulloblastoma (MB) is the most common malignant pediatric brain tumor, characterized by a high propensity for leptomeningeal dissemination. Despite aggressive multimodal therapy, recurrence and metastasis remain the primary causes of mortality. A significant hurdle in improving outcomes is the molecular divergence between the primary posterior fossa tumor and its distal recurrences [1]. Understanding how these cells adapt to diverse neuroanatomical niches requires moving beyond static gene-expression lists toward dynamic regulatory modeling [8]. In this study, we established a novel, minimally invasive microneurosurgical survival platform in mice using the NICO Myriad™ system (Figure 1). By orthotopically grafting pediatric medulloblastoma cells into two distinct neuroanatomical niches, the cerebral cortex (frontal lobe) and the cerebellum, and executing longitudinal resections, we mapped the transcriptomic signatures that regulate local relapse and distant metastasis in cerebellar medulloblastoma. This study uses this novel platform to map the transcriptomic landscape of medulloblastoma, aiming to identify stable, unique targets that persist throughout the tumor’s lifecycle, thereby providing a roadmap for next-generation immunotherapeutic strategies.

**Figure 1:**
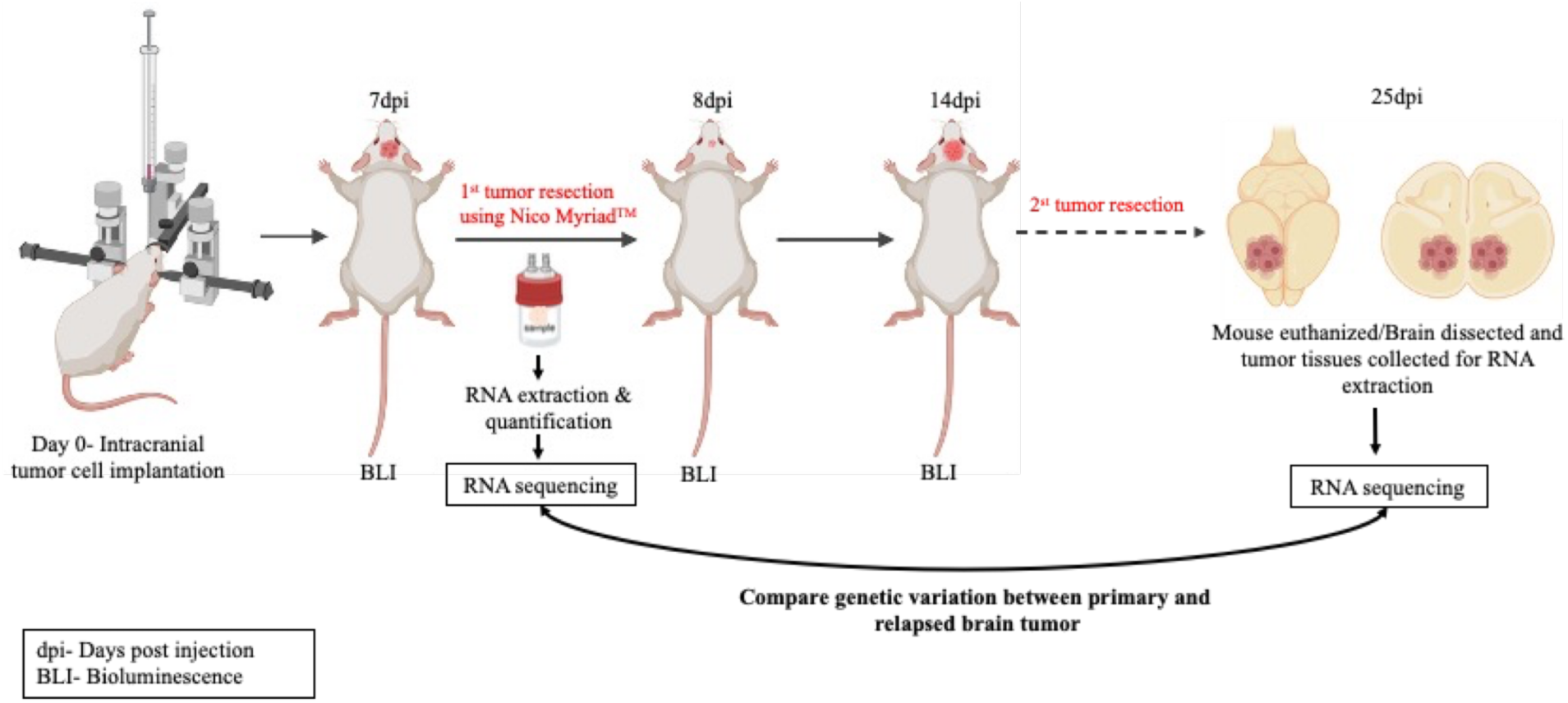
Experimental Schematic of the Novel Microneurosurgical Survival Platform using NICO Myriad™ system showing primary and recurrent tumor resection and its downstream analysis.

## Materials and Methods

### Cell Culture and Reagents

Pediatric medulloblastoma cell line Group III D425 (Group 3/4 characteristics) were cultured in rat-tail collagen (Life Technologies, Cat#A1048301) coated tissue culture plates in glutamine-supplemented medium (Advanced DMEM/F12, 10% fetal bovine serum (Omega Scientific, Cat#FB-02), 1x Glutamax, 1 x Antibiotic-Antimycotic). Cells were maintained in a humidified incubator at 37°C with 5% CO₂. Prior to xenograft, the cells were transduced with a lentiviral vector expressing firefly luciferase and green fluorescent protein (GFP) to enable longitudinal bioluminescence imaging (BLI).

### Animal Studies

This research complies with all the required ethical guidelines and regulations. All protocols related to mouse experiments received approval from the Institutional Animal Care and Use Committee (IACUC) at the University of Southern California (USC). Experiments involving animals were conducted using 7–8-week-old female NOD.Cg-Prkdc^scid^ Il2rg^tm1Wjl^/SzJ mice (The Jackson Laboratory, Cat# 005557, RRID: IMSR_JAX:005557) housed in sterile ventilated cages (5 animals per cage).

### Orthotopic Grafting and Longitudinal Monitoring

Immunodeficient mice (7–8 weeks old) were anesthetized via 2% isoflurane inhalation. Using a stereotaxic frame, medulloblastoma cells (100k cells in 2 µL PBS) were orthotopically implanted into the cerebellum. To model the primary disease, tumor cells were first implanted into the cerebellum at −6mm Bregma, 1mm lateral of the sagittal suture, and 1mm into the cerebellum. The transplanted lines expressed luciferase for *in-vivo* bioluminescent imaging (BLI) post-transplantation. BLI total flux measurements were taken to quantify tumor growth at various intervals, specifically at 3, 7, 14, 16 and 21 dpi. Mice were monitored for presentation of tumor burden related symptoms and humanely euthanized. Tumor progression was monitored through total flux and photos/sec using BLI to establish a baseline for primary tumor growth.

### Microneurosurgical Resection and Survival Model

To recapitulate a clinical recurrence of the disease, a novel two-stage microneurosurgical approach was implemented using NICO Myriad Research Laboratory System (NICO Corporation, USA). This is a minimally invasive technique which standardized to replicate, reduce time and collect tumor samples from preclinical research [9]. Once primary tumors reached BLI threshold 7 dpi, mice underwent a partial or gross total resection of the primary mass. Following tumor resection, a recovery period was maintained to facilitate the survival and subsequent relapse of the malignancy, thereby modeling the longitudinal development of recurrent growth within a living system. To observe if tumor burden has been reduced, BLI was performed the following 24hrs tumor surgical resection and compared. . Mice were monitored daily for recovery and subsequent tumor recurrence. Tumor progression to a terminal stage was characterized by significant tumor relapse, typically occurring at 25 dpi for cerebellar models. Euthanasia was then performed to harvest the recurrent and metastatic tissues for comparative analysis. To harvest the recurrent and metastatic brain tissues with tumor blue polarized light was used to dissect the tumor under a microscope

### Tissue Collection, qPCR and RNA Sequencing

Tissues were harvested at three distinct stages: (1) primary resected tumor (n=7), (2) recurrent tumor at the original site (n=7), and (3) metastatic recurrent tumor (n=7). Tissues were immediately homogenized using the RNeasy Mini Kit (Qiagen) RLT buffer with ß mercaptoethanol in a mortar and a pestle. Further, total RNA from the sample was extracted using the RNeasy Mini Kit (Qiagen) according to the manufacturer’s protocol. RNA quality and concentration was determined using a VarioSkan LUX bioanalyzer (Thermo Scientific) and submitted to Genewiz (USA) for bulk RNA sequencing (n=3). qPCR performed in triplicate per sample as previously described [10]. All primers used for qPCR analysis are human primetime primers purchased from IDT (Integrated DNA Technologies, Iowa, USA).

### Sample QC

Total RNA samples were quantified using Qubit 4.0 Fluorometer (Life Technologies, Carlsbad, CA, USA) and RNA integrity was checked with 4200 TapeStation (Agilent Technologies, Palo Alto, CA, USA).

### Library Preparation and Sequencing

Samples were treated with rRNA depletion using QIAGEN FastSelect rRNA HMR Kit (Qiagen, Germantown, MD, USA), which was conducted following the manufacturer’s protocol. Strand-specific RNA sequencing library was prepared by using NEBNext Ultra II Directional RNA Library Prep Kit for Illumina following manufacturer’s instructions (NEB, Ipswich, MA, USA). Briefly, the enriched RNAs were fragmented for 8 minutes at 94 °C. First strand and second strand cDNA were subsequently synthesized. The second strand of cDNA was marked by incorporating dUTP during the synthesis. cDNA fragments were adenylated at 3’ends, and indexed adapter was ligated to cDNA fragments. Limited cycle PCR was used for library enrichment. The incorporated dUTP in second strand cDNA quenched the amplification of second strand, which helped to preserve the strand specificity. The sequencing library was validated on the Agilent TapeStation (Agilent Technologies, Palo Alto, CA, USA), and quantified by using Qubit 4.0 Fluorometer (ThermoFisher Scientific, Waltham, MA, USA) as well as by quantitative PCR (KAPA Biosystems, Wilmington, MA, USA).

The sequencing libraries were multiplexed and clustered on the flowcell. After clustering, the flowcell was loaded on the Illumina NovaSeq instrument according to manufacturer’s instructions. The samples were sequenced using a 2x150 Pair-End (PE) configuration. The NovaSeq Control Software conducted image analysis and base calling. Raw sequence data (.bcl files) generated by the sequencer were converted into fastq files and de-multiplexed using Illumina’s bcl2fastq 2.20 software. One mismatch was allowed for index sequence identification.

### Transcriptomic and Pathway Analysis

RNA-sequencing data analysis was performed using the Partek Flow® software (v12.1.0, Partek Inc.). Raw sequencing reads were first subjected to pre-alignment quality control to assess base composition and quality scores. Adapter sequences were removed, and low-quality bases were trimmed using the Trim Bases tool within Partek Flow. The processed reads were then aligned to the human [GRCh38] reference genome using the STAR aligner (v2.7.8a) (Splice Transcripts Alignment to a Reference) with default parameters. Following alignment with STAR, the resulting mapped reads were quantified to the genomic annotation model using the Partek E/M algorithm, which facilitates precise counting of both gene and transcript-level features. To ensure robust statistical power and remove low-abundance noise, the raw counts were subjected to a feature filter with a threshold of 1.0. The remaining filtered counts were then normalized using the median ratio method (DESeq2) to account for variations in sequencing depth and library composition across samples. Finally, differential expression analysis was performed using DESeq2 to compare transcriptomic profiles between primary and recurrent tumor tissues.

Statistical significance was defined using an adjusted p-value (P_adj_) < 0.05 and an absolute log2 fold change. Principal Component Analysis (PCA) was performed to visualize transcriptomic variance across cell culture, primary, recurrent, and metastatic stages. Further differential analysis of the samples was performed with FDR step up >= 0.05. Overlapping gene signatures within the cerebral cortex vs. cerebellar regions were identified using Venn diagram analysis to isolate core transcriptomic drivers of the disease. To determine biological significance of these genes, significantly altered genes (fold change >2, p-adj < 0.05) were further analyzed using Ingenuity Pathway Analysis (IPA, Qiagen).

### Functional Enrichment and Subcellular Localization Analysis

Canonical pathway enrichment was performed using QIAGEN Ingenuity Pathway Analysis (IPA). Statistical significance was determined using Fisher’s Exact Test, with a threshold of - log(p-value) > 1.3 (corresponding to a p-value of 0.05). To identify functionally relevant pathways, a downstream z-score filter of z ≥ 2.0 was applied to predict activation or inhibition states. Only pathways with a ratio of >0.1 and containing a minimum of 5 molecules from the dataset were considered for final interpretation. This platform was used to categorize genes by their activation states, identifying significant upregulation or downregulation patterns in primary and recurrent tumor cohorts. Furthermore, the IPA Molecule Activity Predictor (MAP) and subcellular localization tools were employed to map the protein products of these shared genes to their specific cellular compartments, including the cytosol, plasma membrane, nucleus, and extracellular space. This spatial characterization was critical for prioritizing candidates for precision immunotherapy, specifically identifying overexpressed surface antigens suitable for CAR T-cell targeting.

### Statistical Analysis

Bioluminescence data and gene expression levels are presented as mean ± SEM. Statistical significance between groups was determined using one-way ANOVA or Student’s t-test where appropriate. A p-value < 0.05 was considered statistically significant.

## Results

### Validation of In vivo Microneurosurgical Resection and Recurrence in Cerebral cortex and Cerebellum

To evaluate brain tumor relapse following surgical intervention in distinct neuro-anatomical compartments, we established an orthotopic murine model coupled with longitudinal bioluminescence imaging (BLI) and microneurosurgical resection using the NICO Myriad™ system (**Figure 1**). In the cerebral cortex (frontal lobe) model, primary tumors established a bioluminescent signal by 7 days post-injection (dpi) (**Figure 2A**). Following primary tumor resection using the NICO Myriad™ system, BLI total flux showed a statistically significant reduction in tumor burden at 8 dpi (*P=0.0454*, **Figure 2B**). Subsequent longitudinal BLI revealed aggressive local and metastatic tumor recurrence by 15 dpi, with total flux significantly exceeding both post-resection levels at 8 dpi (*P< 0.0001*) and primary pre-resection levels at 7 dpi (*P= 0.0032*, **Figure 2B**). Post-mortem brain shows gross fluorescent tissue confirming tumor recurrence within the primary surgical cavity (white arrowhead) alongside metastatic spread to adjacent cortical areas (yellow arrowhead, **Figure 2C**). To determine whether surgical resection and relapse were consistent in the cerebellum, we performed an orthotopic cerebellar resection model (**Figure 2D**). Primary cerebellar masses grew to a significant volume by 14 dpi before undergoing NICO Myriad™ resection (**Figure 2D**). BLI total flux quantification confirmed significant surgical resection at 15 dpi compared to preoperative baseline at 14 dpi (*P= 0.0019*, **Figure 2E**). By 21 dpi, all animals developed substantial recurrent tumor growth (*P=0.0002*) compared with 15 dpi (**Figure 2E**). Post-mortem analysis of the mouse brain with fluorescent whole-brain sectioning validated primary cavity relapse within the cerebellum (white arrowheads) along with bilateral metastatic dissemination into neighboring cerebellar hemispheres (yellow arrowheads) (**Figure 2F**). Together, these BLI flux differences and histological fluorescence analyses validate the NICO Myriad™ platform as a highly reproducible *in vivo* model for capturing post-surgical local recurrence and distant metastasis across distinct brain compartments.

**Figure 2:**
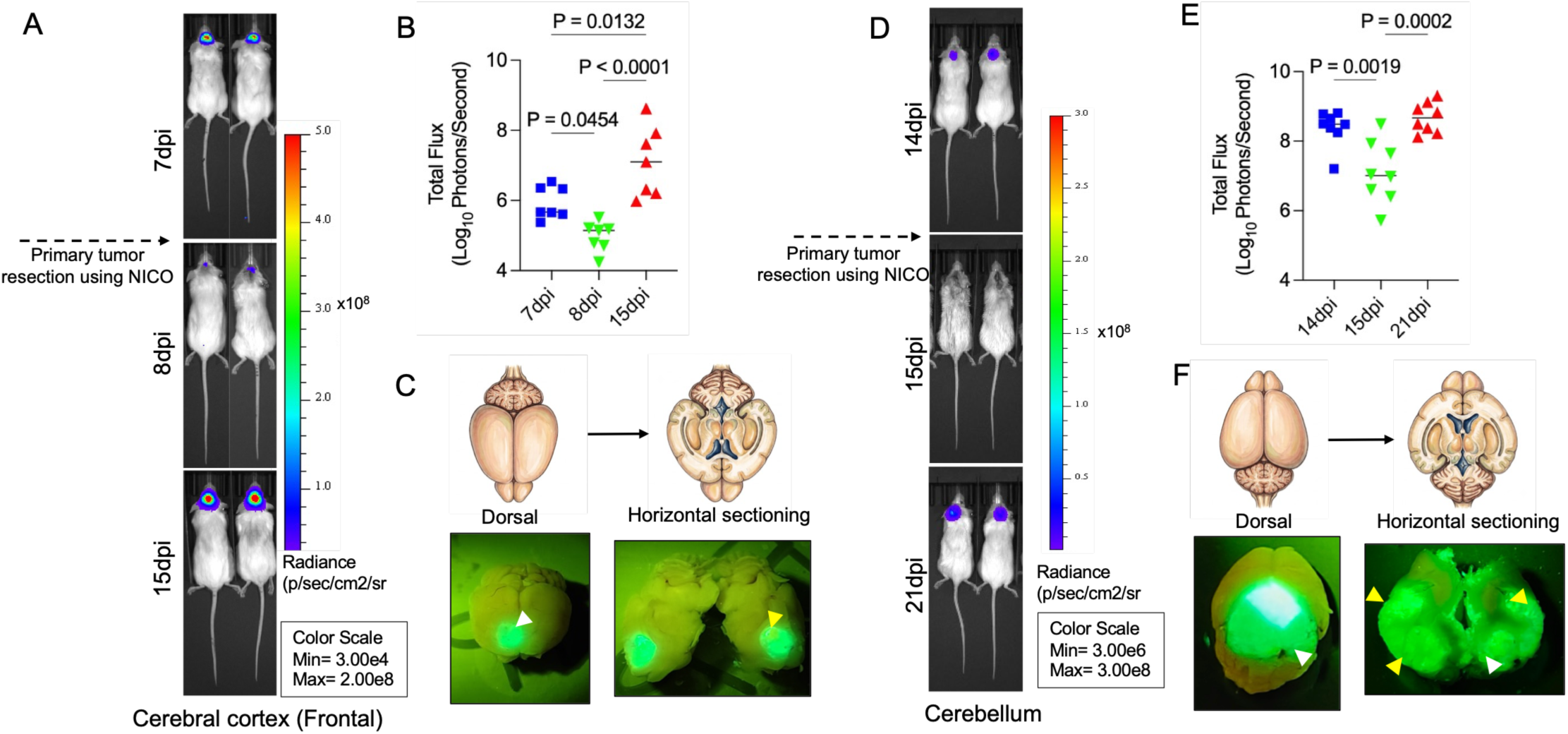
Validation of Microneurosurgical Tumor Resection and Recurrence in Cerebral Cortex and Cerebellar Models. **A-B.** Representative image showing total BLI flux. (log_10_ Photons/Second). in the cerebral cortex cohort and quantification of total BLI flux (log_10_ Photons/Second) in the cerebral cortex cohort showing tumor burden following primary resection (7 dpi vs. 8 dpi, *P = 0.0454*), followed by significant recurrent progression at 15 dpi relative to 8 dpi (*P < 0.0001*) and 7 dpi (*P = 0.0132*)**. C.** Schematic illustration showing ex vivo whole-mount and horizontal sectioning of mouse brain. Representative post-mortem whole-brain fluorescence imaging of the cerebral cortex model highlighted local recurrent primary cavity tumor burden (white arrowhead) alongside distal metastatic dissemination (yellow arrowhead)**. D-E.** Representative image showing longitudinal BLI of orthotopic cerebellar tumors and total BLI flux quantification in the cerebellar model validated substantial surgical debulking between 14 dpi and 15 dpi (*P = 0.0019*), as well as significant recurrent tumor progression by 21 dpi (*P = 0.0002*). **F.** Representative post-mortem fluorescent brain slice images further confirmed primary cavity relapse in the cerebellum (white arrowheads) accompanied by bilateral metastatic dissemination across cerebellar regions (yellow arrowheads). Data points represent individual animals with mean values shown; statistical significance was evaluated using one-way ANOVA with Tukey’s post-hoc test (*P < 0.05*).

### Targeted Gene Expression Profiling Shows Coordinated Transcriptional and Epigenetic Remodeling During Medulloblastoma Relapse

RNA sequencing was used to evaluate how specific molecular networks evolve during post-surgical relapse, quantitative real-time PCR (qPCR) was performed on primary, locally recurrent, and distally metastatic recurrent tissues in both cerebral cortex and cerebellar models (**Figure 3-5**). The common gene candidates were categorized into oncogenes, tumor suppressors, and epigenetic regulators to map tumor stage-specific molecular shifts. Expression of oncogenes show distinct evolutionary transition from primary developmental pathways to post transcriptional mechanisms (**Figure 3, A–L**). *RBM8A* displayed significant upregulation in local and distal recurrent in both cerebral cortex (**Figure 3A**) and cerebellum (**Figure 3G**). Similar observations were seen for *RAB5C* expression (**Figure 3B,H**). *PTCH1,* a Sonic Hedgehog (SHH) pathway component, and *MYCBP2* show significant downregulation in recurrent and metastatic stages across brain regions (**Figure 3C,I)**. *MYCBP2* shows significant downregulation upon relapse (**Figure 3D, J**), whereas *METTL12* (**Figure 3E,K**) and mTORC1 regulator *LAMTOR4* (**Figure 3F,L**), had no change in their expression across all stages. Further, our tumor suppressor gene expression analysis shows a loss of key negative regulators paired with compensatory activation (**Figure 4, A–F**). Canonical tumor suppressor *FOS* (**Figure 4A, D**) showed significant downregulation in local and distal recurrent tumors relative to primary controls. *PTEN* expression showed a significant decrease in the cerebellum; PTEN levels in the cerebral cortex remained unchanged (**Figure 4B,E**). The m6A RNA demethylase *ALKBH5* exhibited significant upregulation in local recurrent and distal tumors across the cerebral cortex (**Figure 4C**) and cerebellar (**Figure 4F**) niches. We performed expression analysis of epigenetic regulators, showing a shift in chromatin accessibility networks (**Figure 5A–F**). *HDAC2* expression was significantly downregulated in local recurrent and distal tumors compared to primary controls in both the cerebral cortex (**Figure 5A**) and the cerebellum (**Figure 5D**). Conversely, *NAA15* showed significant upregulation in the cerebral cortex (**Figure 5B**) in recurrent and metastatic stages, but no change in the cerebellum (**Figure 5E**). Additionally, CDH18 expression in any stage of the tumor at either of the brain regions remains unchanged as well (**Figure 5C, F**). Taken together, the qPCR data demonstrate that medulloblastoma recurrence downregulate primary developmental drivers (*PTCH1*, *MYCBP2*), canonical suppressors (*FOS*, *PTEN*), and chromatin regulators (*HDAC2*), while upregulating post-transcriptional factors (*RBM8A*, *RAB5C*), acetylation regulators (*NAA15*), and m6A RNA modifiers (*ALKBH5*) within the resected brain microenvironment.

**Figure 3.**
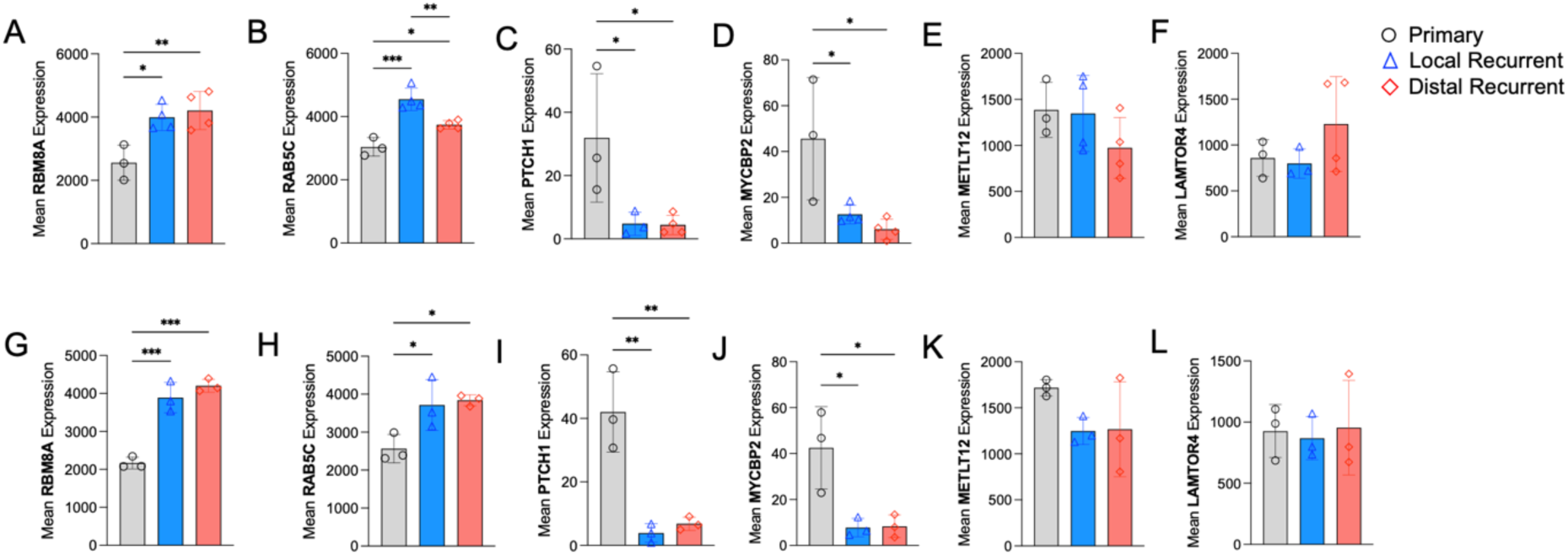
Medulloblastoma recurrence is characterized by downregulation of primary oncogenic drivers and upregulation of vesicular regulators. Quantitative real-time PCR (qPCR) expression profiles of oncogenes across primary (grey), local recurrent (blue), and distal recurrent (red) stages in the cerebral cortex **(Panels A–F)** and cerebellum **(Panels G–L)**. **(A–F)** local and distal recurrent cerebral cortex model show significant upregulation of *RBM8A* (A) and *RAB5C* (B), and significant downregulation of *PTCH1* (C) and *MYCBP2* (D), and no change in *METTL12* (E) and *LAMTOR4* (F) compared to the primary. **(G–L)** Cerebellum model showed similar pattern. There was significant upregulation of *RBM8A* (G) and *RAB5C* (H), downregulation of *PTCH1* (I) and *MYCBP2* (J), and baseline expression of *METTL12* (K) and *LAMTOR4* (L). Bar plots represents mean gene expression ±SEM (n=3). Statistical significance was calculated using one-way ANOVA *(* P < 0.05, ** P < 0.01, *** P < 0.001)*.

**Figure 4.**
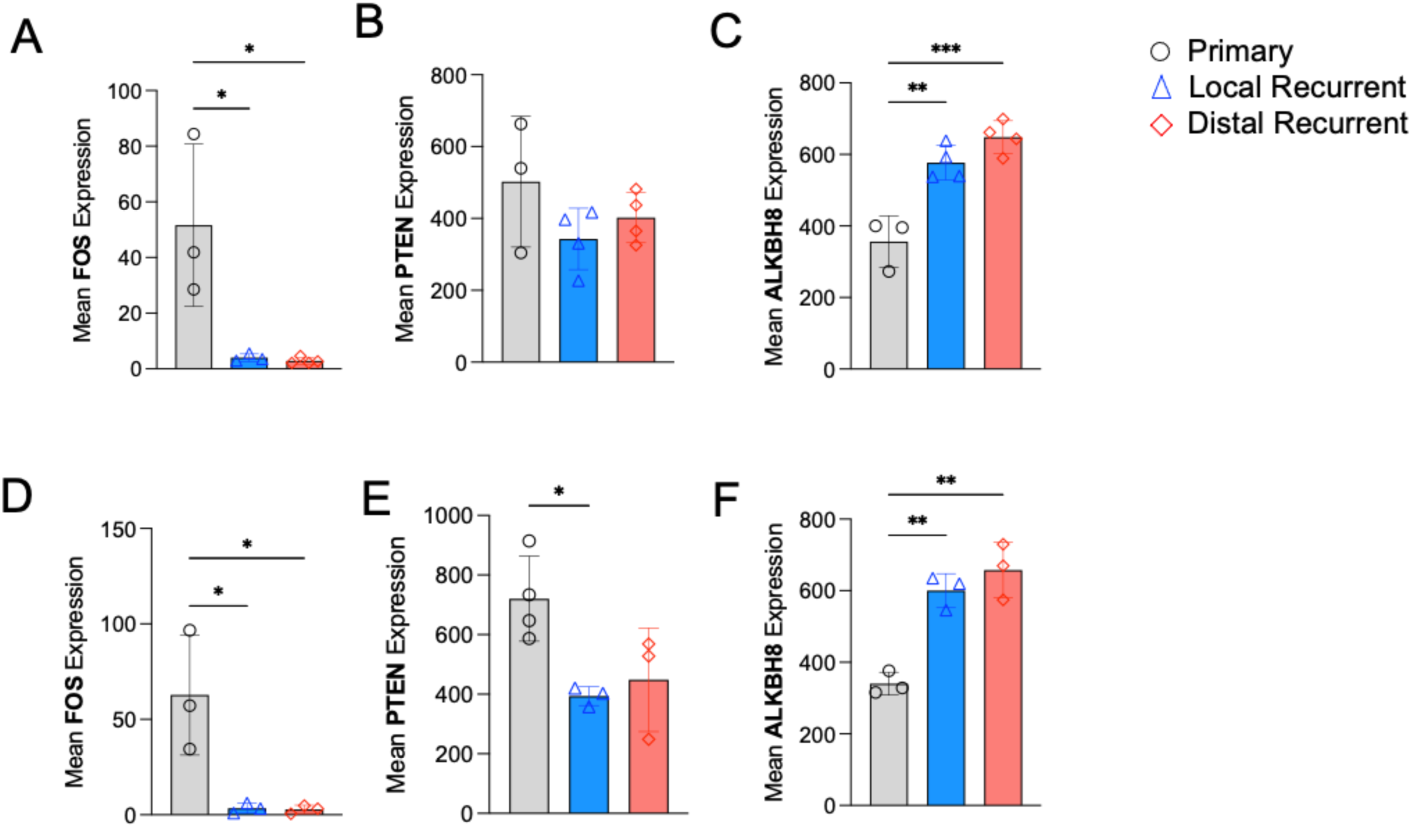
Medulloblastoma recurrence promotes downregulation of canonical tumor suppressors but upregulates RNA demethylation. Quantitative real-time PCR (qPCR) expression of tumor suppressors across primary (grey), recurrent (blue), and metastatic (red) stages in cerebral cortex **(Panels A–C)** and cerebellar **(Panels D–F)** brain regions. **(A–C)** The cerebral cortex shows significant downregulation of *FOS* **(A)** and no change in *PTEN* **(B)**, with significant upregulation of *ALKBH8* **(C)** in local and distal recurrent tumors. **(D–F)** The cerebellar model shows significant downregulation of *FOS* **(D)** and *PTEN* **(E)** and upregulation of *ALKBH8* **(F)**. Data are presented as mean gene expression ±SEM (n=3). Statistical significance was determined using one-way ANOVA *(* P < 0.05, ** P < 0.01, *** P < 0.001)*.

**Figure 5.**
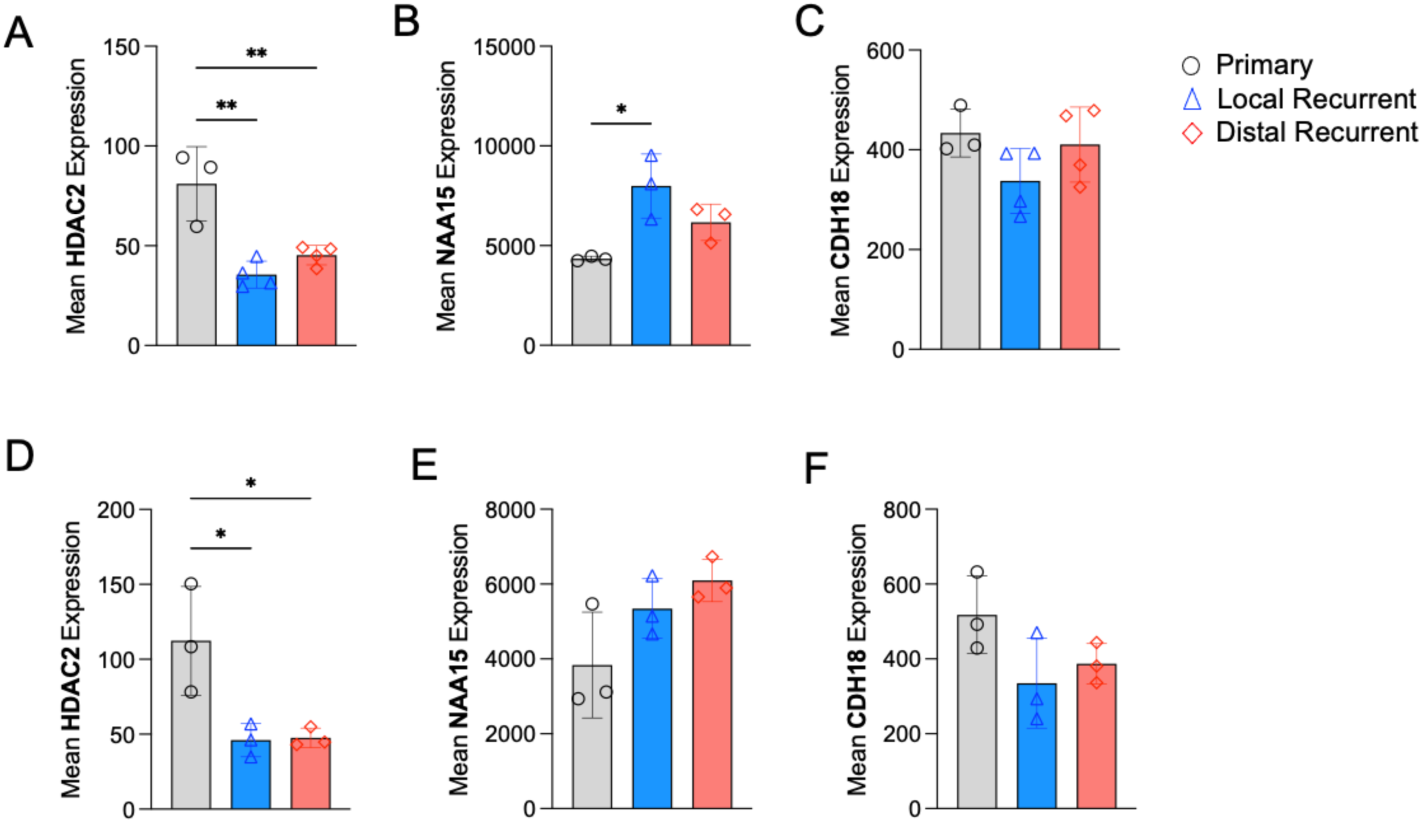
Post-surgical medulloblastoma recurrence alters epigenetic regulators across cerebral cortex and cerebellar niches. Quantitative real-time PCR (qPCR) validation of key epigenetic regulators across disease stages in primary (grey), recurrent (blue), and metastatic (red) tissues. **(A–C)** The cerebral cortex shows significant downregulation of histone deacetylase *HDAC2* **(A)**, upregulation of *NAA15* **(B),** and no change in *CDH18 expression* **(C)**. **(D–F)** Cerebellum local and distal recurrent tumors show significant downregulation of *HDAC2* **(D)**, no change in *NAA15* **(E)** and *CDH18* **(F)** expression. Data presented as mean gene expression ±SEM (n=3). Statistical significance was calculated using one-way ANOVA (*\* P < 0.05, ** P < 0.01)*

### Identification of Conserved Core Gene Networks and Canonical Pathways Across Disease Stages

The transcriptomic profiles of primary, local recurrent, and distal recurrent tumors across cerebral and cerebellar regions show shifts in microenvironmental and stage-specific cellular pathways (**Figure 6**). In the cerebral cortex (frontal lobe), 6,117 unique genes were found in primary tumors, 426 in local recurrences, and 1,105 in distal recurrences; of which 361 genes overlap between local and distal recurrences (**Figure 6A**). Interstage overlap revealed 1,244 shared genes between primary and local recurrent tumors, 1,136 between primary and distal recurrent tumors, and 361 between local and distal recurrences, with a highly conserved core program of 2,702 shared genes across all three stages (**Figure 6A**). Functional annotation of 2,702-gene core using QIAGEN Ingenuity Pathway Analysis (IPA) identified significant enrichment (-log_10_ (*P value*) >1.3) across diverse canonical pathway categories, including cellular growth, proliferation, and development, signal transduction, cellular stress, injury, and intracellular second messenger signaling (**Figure 6B**). Activation z-score revealed positive regulation (orange nodes) across several disease-specific, neural, and growth factor cascades (**Figure 6B**).

**Figure 6.**
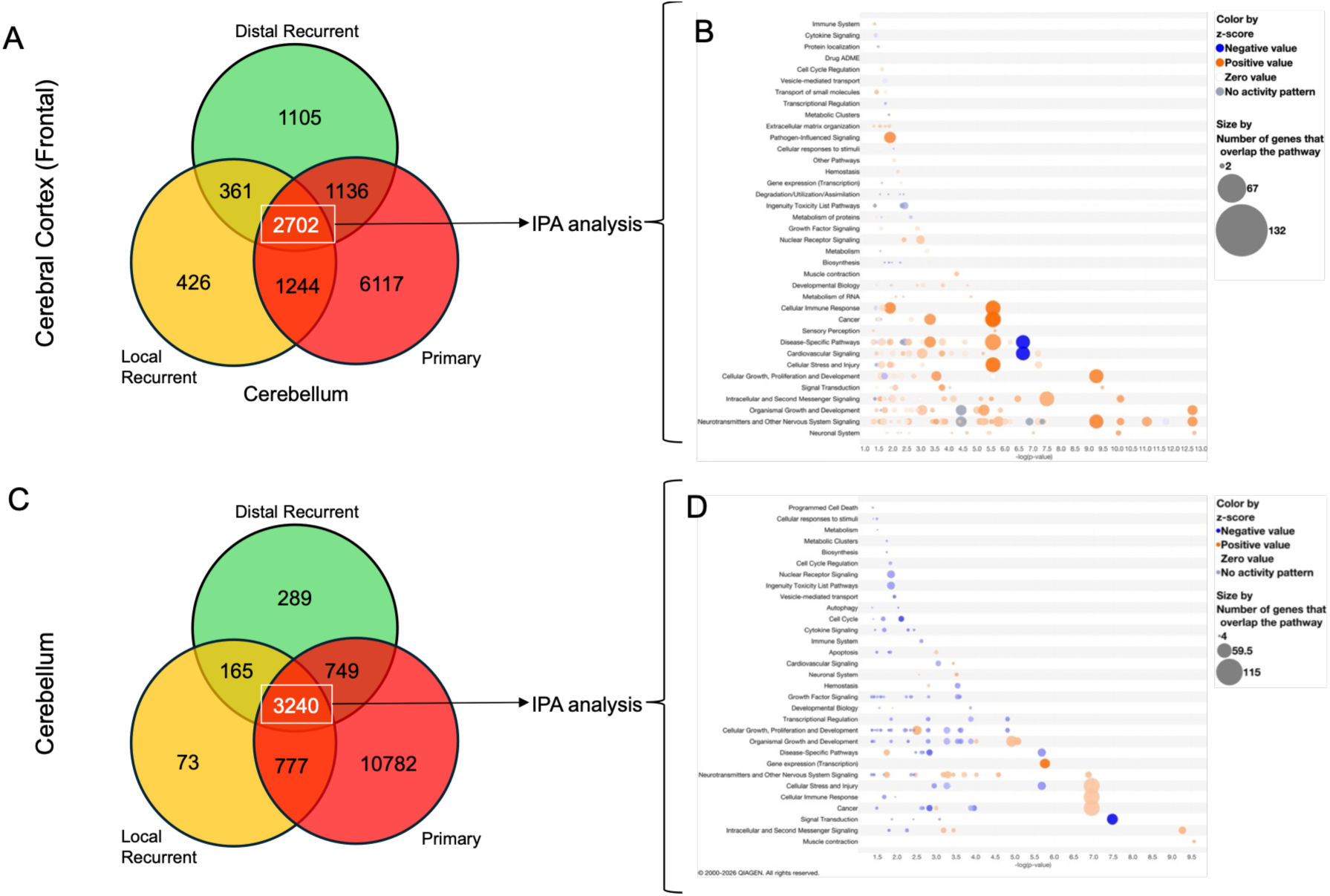
Transcriptomic profiling identifies conserved pathways underlying medulloblastoma relapse in cerebral cortex and cerebellum. **(A)** Venn diagram displaying unique and overlapping gene counts from bulk RNA sequencing across primary, local recurrent, and distal recurrent stages in the cerebral cortex (frontal lobe) model. **(B)** QIAGEN Ingenuity Pathway Analysis (IPA) canonical pathway bubble plot showing enriched functional pathways derived from the 2,702 shared core genes in the frontal lobe model, where bubble size corresponds to the number of overlapping pathway genes (range: 2 to 132 genes) and bubble color indicates the predicted activation z-score (orange = positive activation, blue = negative inhibition, grey = zero/no activity pattern). **(C)** Venn diagram demonstrating transcriptomic intersection across primary, local recurrent, and distal recurrent stages in the cerebellar model. **(D)** QIAGEN IPA canonical pathway bubble plot depicting enriched signaling and metabolic pathways associated with the 3,240 shared core cerebellar genes, with bubble size scaling from 4 to 115 genes and color representing predicted z-score activation states. Statistical significance for pathway enrichment across both models is plotted on the x-axis as – log_10_ (*P*–value).

The cerebellar recurrent model shows an evolutionary architecture distinct from that of the cerebral cortex (**Figure 6C**). Primary cerebellar tumors expressed 10,782 unique genes, whereas local and distal recurrences exhibited 73 and 289 unique genes, respectively (**Figure 6C**). Pairwise overlap included 777 genes shared between primary and local recurrences, 749 between primary and distal recurrences, and 165 between local and distal recurrences (**Figure 6C**). A central core of 3,240 genes showed expression across all three stages in the cerebellum (**Figure 6C**). QIAGEN IPA canonical pathway analysis of these 3,240 genes revealed prominent clustering in cellular immune responses, cellular stress and injury, gene expression/transcription, and cellular growth and proliferation (**Figure 6D**). Together, these bioinformatic analyses demonstrate that medulloblastoma tumors maintain a stable, highly conserved functional backbone of key signaling and metabolic pathways throughout disease progression, yet they also undergo significant transcriptomic adaptation after surgical resection and during metastasis.

## Discussion

The primary clinical barrier in brain tumor management is not the initial response to therapy, but due to divergent recurrence and leptomeningeal metastasis [1]. Traditionally, preclinical models often evaluate primary xenografts or end-stage disseminated disease, which fails to explain the physiological and evolutionary changes of brain tumor recurrence seen post-surgical resection [11]. In this study, by pairing longitudinal resection with stage-specific RNA sequencing and targeted qPCR validation, we demonstrated that post-surgical medulloblastoma relapse is regulated by post-transcriptional RNA processing, transcriptomic adaptation, and chromatin remodeling. A major finding of our targeted qPCR validation is the downregulation of *PTCH1* and *MYCBP2*—in both local surgical cavity relapses and distal metastases. In primary medulloblastoma, particularly Sonic Hedgehog (SHH) subtypes, *PTCH1* mutations drive aberrant precursor proliferation [2, 12]. However, our data show that recurrent and metastatic tumors downregulate *PTCH1*, reflecting distinct primary developmental signatures. This transcriptional shift suggests that therapies targeting classical primary drivers may fail in the recurrent setting due to evolutionary changes [13]. Further, we observed a significant loss in tumor suppressors *FOS* and *PTEN*. Loss of *PTEN* is a well-established driver of aggressive disease, promoting survival and invasive behavior through uninhibited PI3K/Akt/mTOR signaling [14]. *RBM8A* (RNA Binding Motif Protein 8A), a key component of the exon junction complex, and *RAB5C*, a GTPase regulator of endosomal transport, both exhibited progressive, statistically significant upregulation across local and distal recurrences [15, 16]. *RBM8A* has been implicated in driving stemness and aggressive progression in neural malignancies through altered mRNA splicing and translation control [17]. Similarly, upregulation of *RAB5C* expression facilitates endosomal trafficking and cell-matrix interactions necessary for metastasis in brain microenvironments [16, 18]. This highlights a shift toward post-transcriptional adaptive survival mechanisms under the stress of surgical resection. The post-transcriptional adaptation is further accompanied by upregulation of *ALKBH5* across all recurrent stages. As an N^6^-methyladenosine (m^6^A) RNA demethylase, *ALKBH5* regulates mRNA stability and nuclear export, and its overexpression has been linked to glioblastoma stem cell maintenance and metabolic adaptation [19]. At the epigenetic level, we observed a significant downregulation of the class I histone deacetylase *HDAC2*, alongside an upregulation of the acetyltransferase subunit *NAA15*. *HDAC2* loss alters global chromatin accessibility, which can promote genomic instability [20]. In brain tumors, histone deacetylase suppression and altered acetylation kinetics by factors like *NAA15* are frequently associated with enhanced phenotypic plasticity and resistance to cellular stress [21].

While individual gene targets like *PTCH1* are lost during relapse. This indicates that medulloblastoma cells do not stop proliferating upon losing the primary driver; rather, they rewire their signal transduction networks through 2,702 and 3,240 shared gene cores in all tumor stages to sustain growth. Furthermore, the enrichment of cellular stress/injury and immune response pathways in the conserved core reflects the hostile tissue microenvironment encountered after surgery. The physical trauma of microneurosurgical resection triggers local neuroinflammation, tissue remodeling, and hypoxic stress within the resected cavity [22]. Survival within this scar requires tumor cells to upregulate stress-adaptation machinery and inflammatory signaling networks. Additionally, as recurrent tumors undergo extensive clonal selection, targeting transient stage-specific alone can often leads to therapeutic failure [1]. Instead, identifying surface-expressed or enzymatically active candidates within this transcriptomic backbone can provide a roadmap for developing universally applicable CAR T-cell therapies and precision small-molecule inhibitors across all stages of tumor evolution.

In summary, this study establishes a clinically relevant, longitudinal microneurosurgical survival model using the NICO Myriad™ system to track the evolution of medulloblastoma across primary resection, local recurrence, and distant metastasis. Our findings demonstrate that post-surgical relapse is driven by a distinct molecular transformation. Despite these stage-specific expression shifts, bioinformatic profiling identified a highly conserved core transcriptomic backbone across all disease stages and anatomical niches. Targeting this core signature can offer a roadmap for developing next-generation immunotherapies, such as CAR T-cell treatments, capable of preventing relapse and eradicating disseminated medulloblastoma.

## Acknowledgements

We would like to acknowledge patient advocates and all cancer patients, survivors, and family members for their invaluable role in our research. We thank Ivetta Vorobyova for assistance with core services at the University of Southern California. This work was supported by from the Uncle Kory Foundation Fight: Like The Averys F.L.A.G. Grant and NICO Corporation

## References

1. Morrissy, A.S., et al., Divergent clonal selection dominates medulloblastoma at recurrence. Nature, 2016. 529(7586): p. 351–7.

2. Northcott, P.A., et al., Subgroup-specific structural variation across 1,000 medulloblastoma genomes. Nature, 2012. 488(7409): p. 49–56.

3. Martirosian, V., et al., Medulloblastoma initiation and spread: Where neurodevelopment, microenvironment and cancer cross pathways. J Neurosci Res, 2016. 94(12): p. 1511–1519.

4. Ramaswamy, V., et al., Recurrence patterns across medulloblastoma subgroups: an integrated clinical and molecular analysis. Lancet Oncol, 2013. 14(12): p. 1200–7.

5. Yasargil, M.G., A legacy of microneurosurgery: memoirs, lessons, and axioms. Neurosurgery, 1999. 45(5): p. 1025–92.

6. Sanai, N., et al., An extent of resection threshold for newly diagnosed glioblastomas. J Neurosurg, 2011. 115(1): p. 3–8.

7. Weller, M., et al., Rindopepimut with temozolomide for patients with newly diagnosed, EGFRvIII-expressing glioblastoma (ACT IV): a randomised, double-blind, international phase 3 trial. Lancet Oncol, 2017. 18(10): p. 1373–1385.

8. Birk, S., et al., Quantitative characterization of cell niches in spatially resolved omics data. Nat Genet, 2025. 57(4): p. 897–909.

9. Alomari, S., et al., Implementation of Minimally Invasive Brain Tumor Resection in Rodents for High Viability Tissue Collection. J Vis Exp, 2022(183).

10. Das, D., et al., Tumor cells upregulate neurotransmitter GABA in the choroid plexus through STAT6-Bestrophin1 signaling, promoting leptomeningeal dissemination. Neuro Oncol, 2025. 27(6): p. 1476–1490.

11. Wakefield, L., S. Agarwal, and K. Tanner, Preclinical models for drug discovery for metastatic disease. Cell, 2023. 186(8): p. 1792–1813.

12. Goodrich, L.V., et al., Altered neural cell fates and medulloblastoma in mouse patched mutants. Science, 1997. 277(5329): p. 1109–13.

13. Nussinov, R., B.R. Yavuz, and H. Jang, Tumors and their microenvironments: Learning from pediatric brain pathologies. Biochim Biophys Acta Rev Cancer, 2025. 1880(3): p. 189328.

14. Seront, E., et al., PTEN deficiency is associated with reduced sensitivity to mTOR inhibitor in human bladder cancer through the unhampered feedback loop driving PI3K/Akt activation. Br J Cancer, 2013. 109(6): p. 1586–92.

15. Asthana, S., et al., The Exon Junction Complex Factor RBM8A in Glial Fibrillary Acid Protein-Expressing Astrocytes Modulates Locomotion Behaviors. Cells, 2024. 13(6).

16. Reventun, P., et al., RAB5C Increases Endothelial Release of VWF by Regulating Vesicle Trafficking. Arterioscler Thromb Vasc Biol, 2026. 46(3): p. e323000.

17. Lin, Y., et al., RBM8A Promotes Glioblastoma Growth and Invasion Through the Notch/STAT3 Pathway. Front Oncol, 2021. 11: p. 736941.

18. Frittoli, E., et al., A RAB5/RAB4 recycling circuitry induces a proteolytic invasive program and promotes tumor dissemination. J Cell Biol, 2014. 206(2): p. 307–28.

19. Zhang, S., et al., *m(6)A Demethylase ALKBH5 Maintains Tumorigenicity of Glioblastoma Stem-like Cells by Sustaining FOXM1 Expression and Cell Proliferation Program*. Cancer Cell, 2017. 31(4): p. 591–606 e6.

20. Van Bree, B.A. and L.J. Eichner, The impactful role of the HDACs in the regulation of gene expression and as targets for disease therapy. Sci Adv, 2026. 12(23): p. eaee7295.

21. Haase, S., et al., Epigenetic reprogramming in pediatric gliomas: from molecular mechanisms to therapeutic implications. Trends Cancer, 2024. 10(12): p. 1147–1160.

22. Yang, T., R. Velagapudi, and N. Terrando, Neuroinflammation after surgery: from mechanisms to therapeutic targets. Nat Immunol, 2020. 21(11): p. 1319–1326.

